# PAMOGK: A Pathway Graph Kernel based Multi-Omics Clustering Approach for Discovering Cancer Patient Subgroups

**DOI:** 10.1101/834168

**Authors:** Yasin Ilkagan Tepeli, Ali Burak Ünal, Furkan Mustafa Akdemir, Oznur Tastan

**Affiliations:** Faculty of Engineering and Natural Sciences, Sabanci University, 34956, Istanbul, Turkey; Dept of Computer Science, University of Tübingen, 72076, Tübingen, Germany; Dept of Computer Engineering, Bilkent University, 06800, Ankara, Turkey

**Keywords:** Patient Stratification, Graph Kernels, Multi-view Clustering, Pathways

## Abstract

Accurate classification of patients into molecular subgroups is critical for the development of effective therapeutics and for deciphering what drives these subgroups to cancer. The availability of multi-omics data catalogs for large cohorts of cancer patients provides multiple views into the molecular biology of the tumors with unprecedented resolution. We develop PAMOGK (<u>Pa</u>thway based <u>M</u>ulti <u>O</u>mic <u>G</u>raph <u>K</u>ernel clustering) that not only integrates multi-omics patient data with existing biological knowledge on pathways. We develop a novel graph kernel that evaluates patient similarities based on a single molecular alteration type in the context of a pathway. To corroborate multiple views of patients evaluated by hundreds of pathways and molecular alteration combinations, we use multi-view kernel clustering. Applying PAMOGK to kidney renal clear cell carcinoma (KIRC) patients results in four clusters with significantly different survival times (*p*-value = 1.24*e*-11). When we compare PAMOGK to eight other state-of-the-art multi-omics clustering methods, PAMOGK consistently outperforms these in terms of its ability to partition KIRC patients into groups with different survival distributions. The discovered patient subgroups also differ with respect to other clinical parameters such as tumor stage and grade, and primary tumor and metastasis tumor spreads. The pathways identified as important are highly relevant to KIRC. PAMOGK is available at github.com/tastanlab/pamogk

## 1 Introduction

Cancer is a molecularly diverse disease; within the same cancer type, patients bear different molecular alterations, which manifest themselves as different clinical trajectories [6, 58, 28]. Discovering coherent subgroups of patients with similar molecular profiles is essential to developing better diagnostic tools and subtype specific treatment strategies. Knowledge of molecular subtypes is also key to finding the mechanisms that drive these different subtypes to cancer. The problem of stratifying patients based on their molecular profiles is also critical for other complex diseases.

Characterization of patients with omics technologies and the availability of multi-omics data on large cohorts of patients opens up opportunities for better stratification of patients [51, 48, 6]. Towards this goal, several multi-omics clustering methods have been proposed (reviewed in [33]) to integrate the multidimensional data collected on patients. The simple form of integration is early integration. In this case, patient features derived from single omic data are concatenated, and standard clustering is applied to this combined feature representation. However, this approach equally weighs each data type and suffers from a curse of dimensionality as the higher dimensional features dominate the clustering. There are more sophisticated early integration approaches that aim to overcome these problems. iClusterBayes and its earlier variants [41, 26, 25] and LRACluster assume a latent lower-dimensional distribution of data and uses regularization. A second strategy is to deploy late integration approaches [33]. In this case, the samples are clustered with each omic data type separately, and the ensemble’s cluster assignments are combined into a single clustering solution. The consensus clustering by Monti et al. [27] is frequently used for cancer subtyping [13, 51]. PINS[31] and COCA[14] fall into this category as well. These approaches have the drawback that they do not capture the correlations between the different data types. This strategy leads to poor clustering when each view individually contains a weak signal.

Alternatively, several intermediate integration algorithms have been developed [34]. For example, SNF [55] constructs a patient similarity network using each data type, and these similarities are then fused in a single similarity network through an algorithm based on message passing. Meng *et al*. [24] applies dimension reduction to the axes of maximal covariance between data types, JIVE [23] utilizes the variations in data. MCCA [59, 3] extends the canonical correlation analysis (CCA) [11] to a multi-view setting. There are also several algorithms, which are developed as generic multi-view algorithms ([62]). For example, [16] and [4] extend the spectral clustering algorithm [54], which relies on partitioning a similarity network of samples.

There are also kernel-based intermediate integration algorithms, which take a multi-view clustering approach. Kernel methods are powerful methods, where the samples’ similarities are implicitly calculated in a higher-dimensional space [38]. Several generic multi-view kernel clustering methods (reviewed in [61]) have been developed where some have been applied to cancer subtyping [45, 10]. rMKL-LPP [45] extends the multi-view kernel framework [18] to the multi-omics clustering. A kernel matrix is computed from each omic data type, and a linear combination of kernels is sought for the clustering of the patients in kernel k-means. The localized multiple kernel k-means (LMKMM) [10] method also assumes a linear combination of the views but learns a sample specific kernel matrix weight in a k-means framework.

Although corroborating multi-omics data is important to construct a better view of patient similarities, it might not be sufficient to boost the signal as often only a small fraction of molecular alterations is common among the patients. Analyzing molecular data in the context of molecular networks is a widely used approach to overcome this heterogeneity and sparsity problem (reviewed in [5]). In this work, we present PAMOGK, a multi-view kernel clustering approach, which integrates multi-omics patient data with pathways using graphs ^4^. PAMOGK represents each patient as a set of vertex labeled undirected graphs, where each graph represents the gene interactions in a biological pathway, and the vertex labels are assigned based on patient specific molecular alterations. To quantify patient similarity over a pathway and to attain an omic view, we introduce a novel graph kernel, the smoothed shortest path graph kernel (SmSPK), which extends the shortest path graph kernel [2]. While existing graph kernels are designed to capture the topological similarities of the graphs, SmSPK captures the similarities of the vertex label within the graph context. This allows us to capture patients’ similarities that stem from the dysregulation of similar processes in the pathways. By utilizing multi-view kernel clustering approaches, PAMOGK stratifies patients into subgroups. PAMOGK also offers additional insights by showing how informative each pathway and the data type is to the clustering process based on the assigned kernel weights.

We apply our methodology to kidney renal cell carcinoma(KIRC) data made available through the Cancer Genome Atlas Project (TCGA) [29]. We integrate the patient somatic mutations, gene expression levels, and protein expression dataset. Compared to the state-of-the-art multi-omics clustering methods, PAMOGK consistently outperforms in terms of its ability to partition into groups with different prognosis. Extracting of the relative importance of pathways in the clustering process show that the discovered pathways that are relevant to KIRC. PAMOGK is available at https://github.com/tastanlab/pamogk.

## 2 Methods

Given a set of cancer patients, *𝒮*, for which molecular profiles of the tumors are available, PAMOGK aims to stratify them into *k* subgroups through integrating pathways. Formally, we would like to find a partitioning *𝒞* such that: *𝒮* is grouped into *k* number of disjoint subsets *C*_*i*_’s where 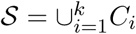 and where *C*_*i*_ ∩ *C*_*i*_ = ∅. In this section, we detail the steps of PAMOGK and data processing used in our experiment. Let *M* be the number of pathways, *D* be the molecular alteration types (mutations, altered expression, etc.) available for the patients and *N* be the number of patients.

### 2.1 PAMOGK Overview

PAMOGK involves three main steps (Figure 1). In the first step, each pathway is represented with an undirected graph. Next, for a given molecular alteration type, i.e., somatic mutations, a patient’s molecular alterations are mapped on the pathway. These alterations constitute the patient-specific node labels of the patient’s graph. Thus, a “view” is constructed for each pathway-molecular alteration type pair. To assess a pair of patients’ similarity under a view, in the second step, the novel graph kernel, SmSPK, is computed to quantify a patient pair’s similarity over a pathway and a molecular alteration type. Each *N × N* kernel matrix constitute a *view* to the patient similarities. In the final step, to stratify cancer patients into meaningful subgroups, these multiple kernels are input to a multi-view kernel clustering algorithm. In the following sections, we elaborate on each step of PAMOGK with more technical details.

**Fig. 1:**
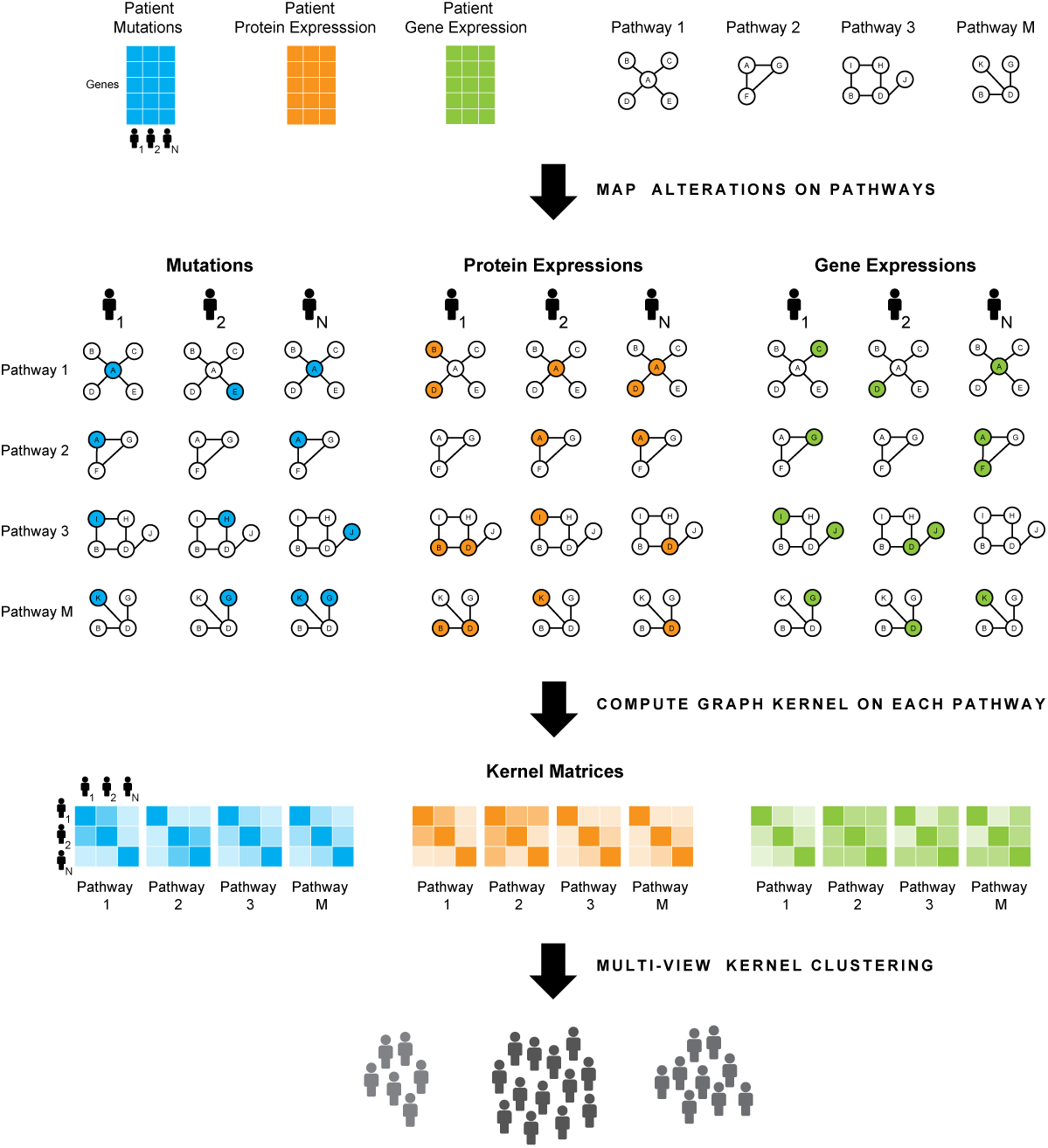
The PAMOGK framework. PAMOGK takes different omic measurements (shown in different colors) and pathways as input. Note that pathway graphs are shown smaller than usual due to size constraints. Each pathway-omic pair constitute a view. In a view, each patient is represented with an undirected graph whose interactions are based on the pathway, and the node labels are molecular alterations of the genes for that patient. For each view, a patient-by-patient graph kernel matrix is computed to assess patient similarities under that pathway-alteration view. In the final step, these views are input to a multi-view kernel clustering method to obtain the patient clusters.

### 2.2 Step 1: Patient graph representation

We first convert each pathway to an undirected graph where nodes are genes, and an edge exists if there is an interaction between the two genes. For each pathway graph *i* and patient *j*, we define an undirected vertex labeled graph 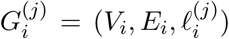. *V*_*i*_ = {*v*_1_, *v*_2_, *…, v*_*n*_} is the ordered set of *n* genes in the pathway *I* and *E*_*i*_ ⊂ *V*_*i*_ × *V*_*i*_ is a set of undirected edges between the genes in this pathway. The label 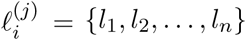 is in the same order of *V*_*i*_ and represents the corresponding vertex’s label for patient *j*. For a specific pathway, the pathway graph structure is the same for all patients and is defined by the set of interactions in the pathway while the vertex labels are different and are based on each patient’s molecular alterations. For a patient *j*, 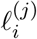 entries are assigned based on the patient’s molecular alteration profile. For example, in the case of somatic mutations, if the corresponding gene *k* is mutated in patient *j*, label of value 1 is assigned to this gene (node), and 0 otherwise. At the end of this step, we have *N* × *M* × *D* labeled pathway graphs.

### 2.3 Step 2: Computing Multi-View Kernels with Graph Kernels

In this step, we would like to assess the similarities of the patients on a given pathway for a given molecular data type. For this, we resort to graph kernel functions. While typical kernels take vectors as input, a graph kernel takes two graphs as input and returns a real-valued number that quantifies the similarity of two input graphs: 𝒦: 𝒢 × 𝒢 ↦ ℝ [52]. Powerful graph kernels are presented in earlier work [42, 2, 30]. However, these graph kernels are designed to compare graphs with different graph structures and to identify similarities and differences that arise from these different structures. In our case, though, we would like to compare graphs with identical topology but different node label distribution. The graphs’ structures are identical because they are from the same pathway, and the label distributions are different because of the patient specific alterations. To assess the similarity of topologically identical graphs with different node label distribution, we devise a new graph kernel for our purposes.

Inspired from the shortest path graph kernel [2], SmSPK makes use of all shortest paths of the graphs to characterize them. We also smooth the node labels of a patient in the pathway so that if two patients have alterations in genes in close proximity, they contribute to the similarity even though the set of altered genes are not identical. To propagate node labels along the pathway, we use the random walk with restart, which is a common strategy used in various tasks (reviewed in [5]). For a single graph indexed by *g*, the label propagation is performed by employing the following formula for all patients:

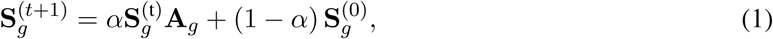

where **S**_**g**_^(0)^ is a patient-by-gene matrix which represents the labels of the vertices in the graph *g* at time *t* = 0 and each row (patient) is determined by 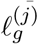 In this case, 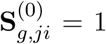 where *j* is index of the patient and *i* is index of the vertex. **S**_**g**_^(*t*)^ is the node label matrix at time *t*. **A**_*g*_ is the degree normalized adjacency matrix of the pathway graph *g. α ∈* [0, 1] is the parameter that defines the degree of smoothing. We iterate over propagation until convergence is attained. We assign node attributes of the graph for each patient based on the final **S**. Once we attain the label smoothed graphs of 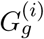 and 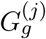, we compute the similarities of these two graphs to each other as follows:

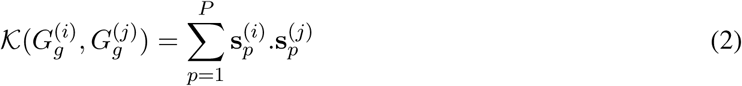

Here, 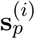 is the vector that represents the labels of the vertices of the graph *G*_*g*_ on the shortest path *p* for patient *i* after smoothing, *P P* is the number of all pairs of shortest paths on the graph. The above function is a valid kernel function, as the dot product is the linear kernel, and the kernel property is preserved under summation.

For a given molecular alteration type and a pathway, we compute the kernel over all pairs of patients. **K** matrix is a symmetric *N* × *N* matrix, for which the *i, j*-th entry is the kernel function evaluated for patient *i* and patient *j* pair. By computing kernel matrices for each pathway and for each molecular alteration type, **2.4** we obtain *M* × *D* kernel matrices. We normalize the /kernel matrices such that all kernel entries are in [0 − 1] by dividing the kernel matrix entry **K**(*i, j*) entry by 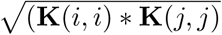

### 2.4 Step 3: Multi-View Kernel Clustering

Each of the kernel matrices computed in the previous section represents a view of the patients’ similarities. To integrate these views, we resort to existing multi-view kernel clustering approaches. We experiment with different approaches (see Section 3.3); multiple kernel k-means with matrix-induced regularization (MKKM-MR) [21] performs the best. Thus the final model of PAMOGK uses MKKM-MR; yet, this step can be replaced by any multi-view clustering approach as long as the method accepts kernel matrices as input. In this section, for completeness, we provide a brief overview of the selected multi-view kernel clustering methods with which we experimented.

#### Multiple Kernel K-Means with Matrix-Induced Regularization

MKKM-MR algorithm objective is to minimize sum-of-squared loss over the cluster assignments. To reduce redundancy among kernel matrices and enhance the diversity of the selected kernel matrices, MKKM-MR [21] uses the matrix-induced regularization. The algorithm solves the following optimization problem:

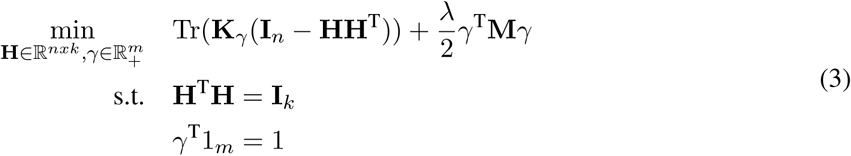

Here, *k* is the number of clusters, *n* denotes the number of samples, *m* is the number of kernel matrices. **H** is the relaxed clustering assignment matrix, *γ* = [*γ*_1_, *γ*_2_,, *γ*_*m*_] are the weights of input kernel matrices. **K**_*γ*_ is the best kernel matrix, **M** is the matrix that measures the relation between kernel matrices. **I**_*x*_ is the *x*-by-*x* dimensional identity matrix, 1_*m*_ is *m* dimensional vector of ones. *λ* is the parameter that adjusts the trade-off between clustering cost and the regularization term.

#### Average Kernel K-Means (AKKM)

Kernel k-means (KKM) [37] is simple but a strong baseline. Since it accepts a single kernel matrix, we input the average of the kernel matrices. We will refer to this method as average kernel k-means (AKKM).

#### Localized multiple kernel k-means (LMKMM)

LMKMM [10] is another powerful method that optimizes not only the weight of the kernel matrices but also the weight of the samples. We reimplemented LMKKM in Python, which is originally provided in Matlab and R.

#### Similarity Network Fusion (SNF)

SNF[56] calculates a similarity matrix of samples using an exponential kernel based on the view created by each data types separately, and construct a similarity network for each view. In these networks, samples are nodes and edge weights are the similarities. Through an iterative procedure based on message passing, the networks are fused into a single network. We use this fusion step of SNF as a multi-view clustering method, where we compute the similarities based on smSPK and cluster the samples with kernel k-Means or spectral clustering.

### 2.5 Dataset and Data Preprocessing

#### Pathway data

As the pathway source, we use National Cancer Institute – Pathway Interaction Database (NCI-PID) at NDEXBio [36]^5^. NCI-PID is a curated database with focus on processes that are relevant to cancer research (download date: Apr 24, 2019). We filter out a pathway if it does not contain any overlapping gene with the omic data genes, which leaves out 165 pathways. The statistics of pathways sizes are provided in Supplementary Table 2.

#### Patient molecular and clinical data

The molecular and clinical data for KIRC is obtained from the TCGA PanCancer project [57]. We retrieve the data directly from Synapse^6^. We only consider the primary solid tumor samples and make use of three different molecular data types that can directly be mapped to pathways: somatic mutations, transcriptomics, and proteomics data. The transcriptomic data include the RNAseq gene expression levels, while protein expression is quantified through Reverse Phase Protein Array (RPPA). The exact data files are listed in Supplementary Table 1.

**Table 1:** The runtimes in seconds for clustering 361 KIRC patients with the three types of omic data for different methods and PAMOGK.

| Method | PAMOGK | LRACluster | PINS | SNF | rMKL-LPP | iClusterBayes | Spectral | K-means | MCCA |
| --- | --- | --- | --- | --- | --- | --- | --- | --- | --- |
| Time | 352 | 289 | 56 | 7 | 109 | 10,898 | 3 | 47 | 6 |

**Table 2:**
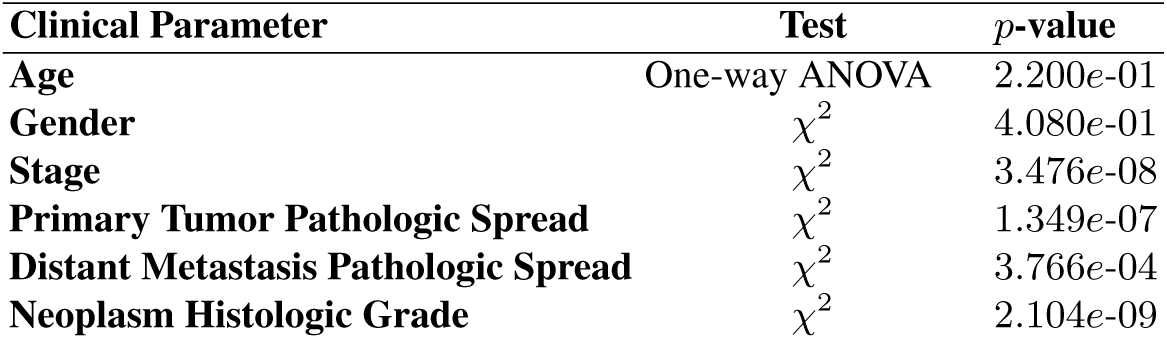
Summary of statistical analyses of clinical variables.

#### Assigning node labels based on molecular alterations

In the case of mutations, the patient node label is assigned as a binary label based on the presence or absence of the mutation. In the expression datasets, the gene and protein expression values are normalized and converted to z-scores relative to other patients. For each data type, if a gene(a protein)’s z-score is greater than − 1.96(which stands for 95% confidence), the gene(the protein) is considered overexpressed while the genes(the proteins) with z-score lower than 1.96 is considered underexpressed. For the underexpressed and the overexpressed values, we use the *z* value as the node attribute. If the z-values are between −1.96 *< z <* 1.96, the node label is assigned to zero. In assigning the expression values, we considered different alternative methods; as discussed in the results section, yields slightly better results. Finally, from the three different omic data sources, five different types of alterations are defined: absence and presence of a somatic mutation in a gene, over and underexpression based on gene expression, over and underexpression based on protein expression.

## 3 Results and Discussion

### 3.1 Experimental Set up

We apply PAMOGK to discover different subgroups of KIRC patients. The dataset contains 361 patients whose molecular profiles come from three separate data types: somatic mutation, gene expression, and protein expression. We define five different molecular alteration types based on these three types of omics data (see Section 2.5). We compute one kernel matrix for each pathway-molecular alteration type; this results in 825 kernels (165 pathways × 5 molecular alteration types), each one of which constitutes a distinct view.

Throughout all experiments, we evaluate four different cluster numbers, *k* = 2, 3, 4, 5. When computing SmSPK, we try 12 different alpha *α* values (Supplementary Table 3). We conduct experiments by using different multi-view clustering methods. These include average kernel k-means(AKKM), LMKKM, MKKM-MR, SNF-kernel k-means and SNF-spectral clustering. If a pathway kernel includes a few or no altered genes, we eliminate it before inputting it into multi-view kernel clustering methods to increase time efficiency. The criteria for this is to eliminate those whose nonzero entries constitute at most 1% of all entries. The parameter *λ* in MKKM-MR is chosen using grid-search. (Supplementary Table 3).

We evaluate the clustering solutions through survival analysis in accordance with previous work [20, 17, 35, 9]. We compare the survival distributions of the clusters using Kaplan-Meier (KM) survival curves [15] and log-rank test’s *p*-value [12]. In the log-rank test, we test whether there is a statistical difference between the survival times of the clusters. In comparing alternative methods, we use the *p*-value of this log-rank test as the performance criteria.

### 3.2 Assessing the Need of a new Graph Kernel

Constructing kernels, which reflect the similarity of patients, is a crucial step of PAMOGK. First, we would like to understand whether there is any merit in using SmSPK as opposed to deploying an already existing and powerful graph kernel. The motivation behind proposing a new kernel is that the existing graph kernels are designed to capture topological similarities. Since we compare the two patients on the same pathway, the structure of graphs shall always be the same. On the other hand, the node label distribution is different as it is patient specific. Thus, the existing graph kernels computed over the same pathway will consider patients as overly similar and would not serve our purpose. To check if this intuition holds, we analyze the distribution of the kernel values computed over all the pathways and the overexpressed molecular alteration type. Since the overexpressed genes are the densest kernels, we choose this data type. We compare SmSPK with kernels that accept continuous node attributes. These kernels include the propagation kernel [30], Graph Hopper kernel [8] and Wasserstein Weisfeiller Lehman graph kernel [47]. We use the implementation provided by the Grakel library [43] for propagation and graph hopper kernel. We use WWL Python library[47] provided by authors for the Wasserstein Weisfeiler Lehman graph kernel. We also tried to compare our method with node attributed shortest path graph kernel [2], but due to slow run times, we abandoned this comparison. Also, to asses, whether we need a graph kernel at all, we also compare to the case where RBF kernel is used. When RBF kernels are computed, they are directly evaluated on the omic data. Thus, they are computed over all the genes regardless of their participation in a pathway. The gamma values of RBF is determined by the median heuristic [39] (Supplementary Table 3). In order to make the comparison fair, we also apply smoothing and choose the results with the best smoothing parameter assignment for each method (Supplementary Table 3).

First we analyze the kernel values computed by each kernel. Figure 2a display the heatmaps computed by each kernel method on an example pathway (more examples are provided in Supplementary Figure Figure 1). The rows and the columns are patients the cell entries colors are proportional to the kernel value computed for the two patients over a single pathway and data type. Figure 1 clearly shows how kernels assign patient similarities of 1 very frequently. To better analyze this over all pathways, we analyze the distribution of kernel values assigned to patients by each of the different kernels. We bin the kernel matrix entries into groups for each kernel and calculate the frequency of each bin. Next, we calculate the average frequency for each bin across all kernel matrices computed (Figure 2b). Figure 2b shows, for each graph kernel, how the kernel values are distributed on average. All the kernels other than SmSPK, assign patient similarities of 1 very frequently (the darkest bin). These results confirm our intuition that due to the identical graph structures, the existing graph kernels are unable to distinguish patients with different molecular alterations on the same pathway graph.

**Fig. 2:**
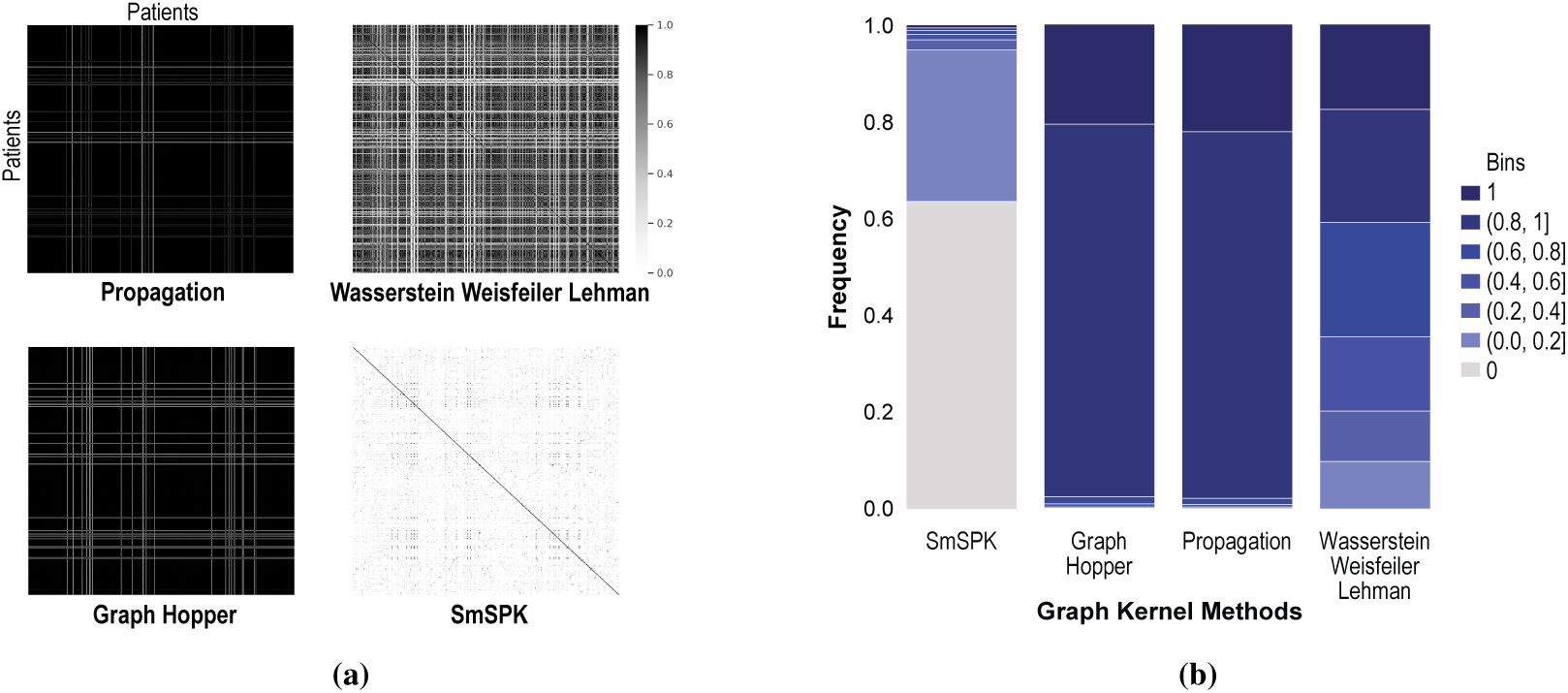
**a)** Example heatmaps of patient-by-patient kernel matrices calculated by different kernel choices. The kernel functions include the propagation kernel, graph hopper kernel, Wasserstein Weisfeiler Lehman, and SmSPK graph kernel methods. Each kernel belongs to *the direct p53 effectors pathway* and overexpressed gene data type. The color black indicates that the similarity of the two patients is evaluated as 1. **b)** The frequency of patient similarities for different kernels over all pathways with the overexpression molecular data. For example, the darkest navy indicates the kernel value of 1, and the height of this bar is the proportion of patient-pairs for which the kernel value is 1. All the kernels other than SmSPK assign patient similarities of 1 very frequently.

Additionally, we compare performances of the kernels based on the lowest *p*-value attained in the logrank test on the survival distributions of clusters. In each experiment, each kernel is used with MKKM-MR method, and they are allowed to choose from a set of predetermined values for each of the hyperparameters. These include *k* for clustering, the smoothing parameter *α* for SmSPK, *λ* for MKKM-MR. The best clustering solution obtained for each method is compared in Figure 3a. We observe that SmSPK outperforms other graph kernels compared. This can be explained based on the previous remark that this graph kernel is formulated to distinguish graphs with different topologies. Additionally, although the use of RBF kernel generally yields good results, the integration of pathway information through SmSPK brings an improvement to the cluster separations in terms of survival.

**Fig. 3:**
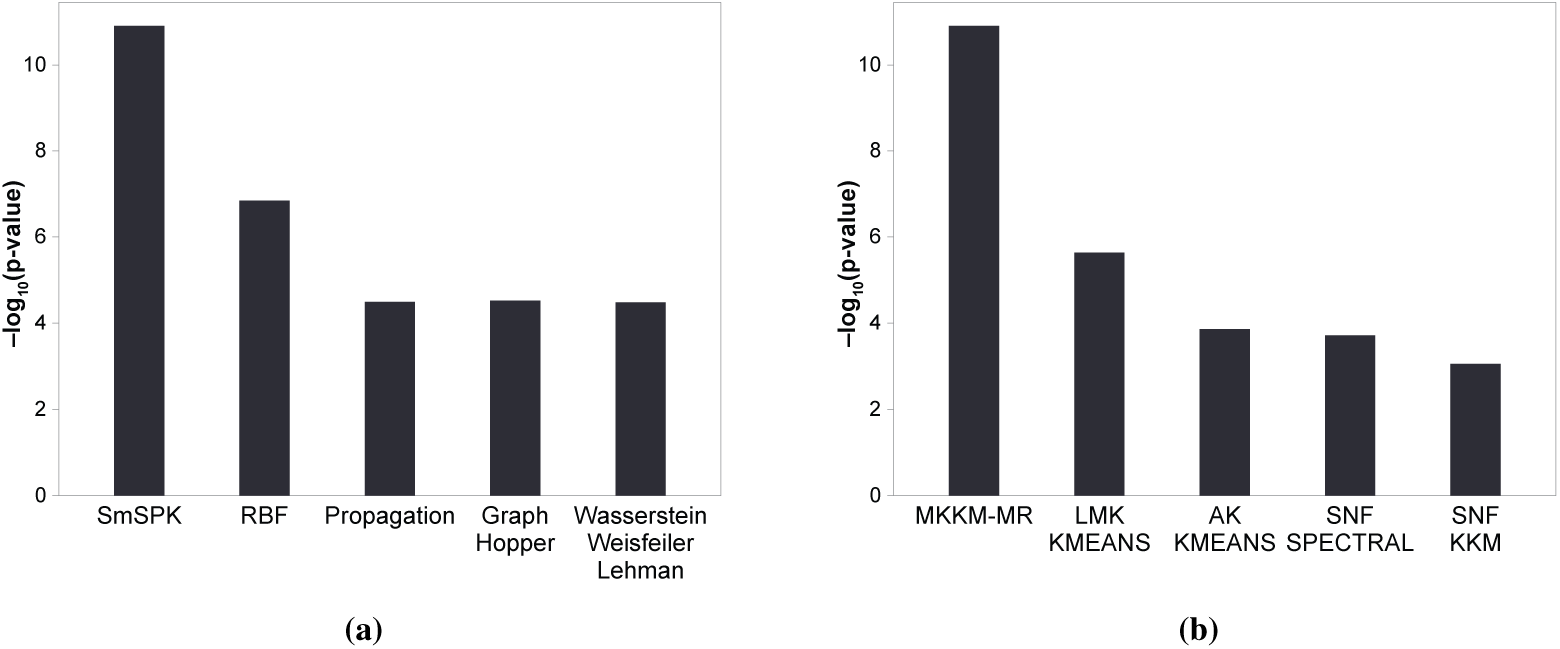
**a)** The log-rank test *p*-values obtained with different choices of kernels employed with MKKM-MR multi-view kernel clustering algorithm. Kernel construction methods consist of SmSPK(our method), propagation graph kernel [30], graph hopper kernel [8], Wasserstein Weisfeiler Lehman graph kernel [46] and radial basis function (RBF) kernel. **b)** The log-rank test *p*-values obtained with different choices of multi-view kernel clustering methods withSmSPK as the kernel construction method. The clustering methods include average kernel k-means (AKKM), localized multiple kernel k-means (LMKKM) [10], multiple kernel k-means with matrix-induced regularization (MKKM-MR) [21], SNF[55] with spectral clustering and kernel k-means(KKM).

### 3.3 Deciding on the Multi-view Kernel Clustering Algorithm to Use in PAMOGK

To determine the multi-view kernel clustering algorithm to be used in PAMOGK, we experiment with different alternatives. The multi-view kernel clustering methods that we analyze include the MKKM-MR [21], AKKM, LMKKM [10] and SNF [55] with KKM(Kernel K-Means) and spectral clustering (see Section 2.4. For each method, we report the best clustering solution, which is determined based on the lowest *p*-value attained in the log-rank test on the survival distributions of clusters. In each experiment, we allow the methods to choose from a set of predetermined values for each of the hyperparameters. These include *k* for clustering, the smoothing parameter *α* for SmSPK, *λ* for MKKM-MR.

Figure 3b summarizes the results in these experiments for the best clustering solution, where *k* = 4. When comparing the three multi-view kernel clustering methods, we observe that MKKM-MR produces the best results. LMKKM, AKKM and SNF based methods yield similar results with the difference that LMKKM performs slightly better than others. Overall, PAMOGK that uses the MKKM-MR multi-view clustering outperforms all the other clustering alternatives. Thus, we employ MKKM-MR in PAMOGK.

The best clustering solution by PAMOGK is obtained when *k* = 4, smoothing parameter, *α* is set to 0.3, and *λ* for MKKM-MR is set to 8. The KM plot of the resulting clustering is provided in Figure 4a. The survival distributions significantly differ (log-rank test, *p*-value = 1.24*e*-11). We should note that the solution with *k* = 3 is also quite good, *p*-value = 8.13*e*-11 (Supplementary Figure 4b).

**Fig. 4:**
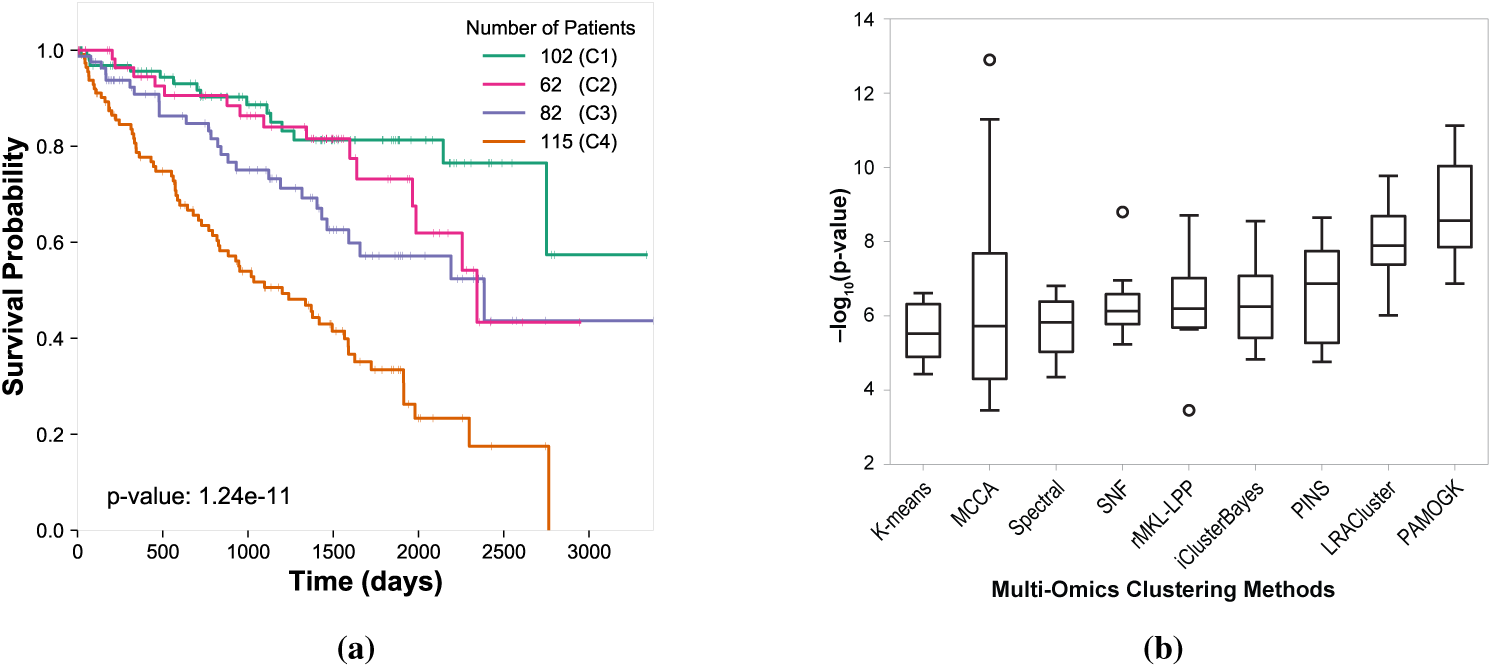
**a)** Kaplan-Meier survival curves of the best clustering solution for KIRC. The *p*-value was obtained from a log-rank test between the groups **b)** Comparison of PAMOGK with the multi-omics clustering methods over 10 different trials. Each trial contains a random subsample of KIRC patients. The boxplot shows the −log (*p*-values) of the log-rank tests conducted on the survival distributions of these clusters. The higher the values, the better the clusters are separated in terms of survival distributions. (Note that PINS method results are over 9 experiments since in one of trial, it did not return a result.)

### 3.4 Effect of Other Design Choices

We checked some of the design choices. The first one is that the alternative strategies of how to map the expression changes on the network. We compare the following three alternatives. In the first one, we use the z-score as the node attribute. In the second one, we assigned the node attribute a binary value (0 or 1) depending on it is overexpressed or not, and similarly for the underexpressed case, underexpressed or not. The third one is the case; we assigned the z-scores of expression if the gene (or the protein) is over or underexpresssed to the degree of the change, and is 0 if it is not over or underexpressed. When we compare these three alternatives, we observe that the directly mapping z-scores do not yield good results, probably because when network smoothing is applied, everything accumulates change. The binary scoring yields also good results, but the third alternative yields better results. This comparison is shown in Supplementary Figure 2. We also assessed whether smoothing effects or not. Supplementary Figure 3 shows that smoothing helps, especially with mutation data, as these alterations are sparse compared to expression data.

### 3.5 Comparison with the State-of-the Art Multi-Omics Methods Performance comparison

We compare PAMOGK with eight other multi-omics methods. These include k-means [22], MCCA [59], LRACluster [60], rMKL-LPP [45], iClusterBayes [25], PINS [31], SNF [55], and finally Spectral Clustering [63]. These methods cover all methods that are included in a recent comparative benchmark study by Rappoport et al. [34] with the exception of multiNMF [19], which we are not able to run properly. In running these algorithms, we set the maximum number of clusters to five and choose the other parameter configurations for each algorithm exactly as in the benchmark study [34].

To assess the performance of different methods, we repeatedly subsample the original patient set, and for each subsample, run the algorithms to find the patient clusters. Each subsample contained 300 patients. Due to prohibiting runtime of iClusterBayes, we were able to conduct this experiment 10 times. The distribution of log-rank test *p*-values attained by each method is displayed in Figure 4b. The comparison over ten runs shows that PAMOGK is the best performer among the nine methods. Not only the median performance is high, but even the 90-th percentile of the trials is superior to almost all methods. It also displays low variance across different runs. For all methods, for all trials, the resulting clusters are balanced in terms of the number of patients participating in the clusters except two trials of MCCA. The log-rank test is known to result in unrealistically low *p*-values when one of cluster size is small [50]. In those two trials, MCCA’s extremely low *p*-values are due to cluster sizes of 9 and 14.

### Runtime comparisons

We conduct a runtime comparison of the algorithms for clustering all the KIRC patients using the three different data types. PAMOGK demands more time to run in comparison to the other methods, with the exception of iClusterBayes. This is because it calculates many more views of the data based on pathways. A second time limiting step is the weight optimization of the kernels in the MKKM-MR algorithm. Despite these additional requirements, the runtime is within reasonable limits, and a typical run takes less than 30 minutes without any parallelization. Replacing the multi-view clustering step with a less demanding algorithm and parallelization could reduce the runtime. Experiments are conducted on the following system configuration: CPU: Intel(R) Xeon(R) CPU E5-2640 v4 @ 2.40GHz CPU. Memory: 256Gb. Operating system: Ubuntu 16.04.4 LTS.

### 3.6 Detailed Analysis of KIRC Subgroups Discovered by PAMOGK

#### KIRC Subgroups’ Associations with Other Clinical Parameters

We analyze the association of clinical parameters of the discovered subgroups other than survival. The parameters include age, gender, tumor stage, primary tumor pathological spread, distant metastasis pathological spread and neoplasm histological grade. The associations of categorical variables are determined using *χ*^2^ test while the continuous variables are tested with one-way ANOVA. We find no statistically significant difference in terms of age (*p*-value = 0.220) and gender (*p*-value = 0.408). All the other clinical parameters differ across groups at a statistically significant level (see Table 2). The distributions of these variables across groups are provided in Supplementary Section 3. The best prognosis group is cluster 1, and the worst prognosis group is cluster 4. For all prognostic tumor-related features, cluster 1 always has more patients with a lower degree stage and grade, whereas cluster 4 always has more patients with a higher degree stage and grade. Overall, this analysis provides additional evidence that PAMOGK partitions KIRC patients into clinically meaningful subgroups The best prognosis group is cluster 1, and the worst prognosis group is cluster 4 (Figure 4a). There are clear differences between these two groups in terms of these additional clinical parameters. More specifically, 53.9% of the patients in cluster 1 are in stage I, whereas 67.8% of the patients in Cluster 4 are either in stage III or Stage IV. (Supplementary Table 6). Also, nearly half of the patients in cluster 1 have primary tumor T1, whereas 60% of the patients in cluster 4 have primary tumor T3 (see Supplementary Table 7). While only 8.82% of the patients of cluster 1 have distant metastasis, this ratio is 29.6% for cluster 4 patients. (Supplementary Table 8). Finally, the fraction of cluster 1 patients with histologic grade G1 and G2 is 61.7%, and those with G4 is 5.9%. For cluster 4, the percentage for G1 and G2 drops to 20.5% and G4 increases to 35.7%. (Supplementary Table 9). For all prognostic tumor-related features, cluster 1 always has more patients with a lower degree stage and grade, whereas cluster 4 always has more patients with a higher degree stage and grade. Overall, this analysis provides additional evidence that PAMOGK partitions KIRC patients into clinically meaningful subgroups.

#### Influential pathways and data types

By inspecting the assigned kernel weights, we can quantify the relative importance of pathways and molecular data types. For KIRC (*k* = 4), the *direct p53 effectors* pathway and gene overexpression kernel emerge as the most important pathway-molecular alteration pair (see Supplementary Figure 5 for the top 10 pairs). By averaging the weights associated with each omic data type, we find that the gene expression is the top important data type, while protein expression data have almost no effect on the clustering (Supplementary Figure 6). This could be arising from the fact that the protein expression data covers only a small number of proteins.

**Fig. 5:**
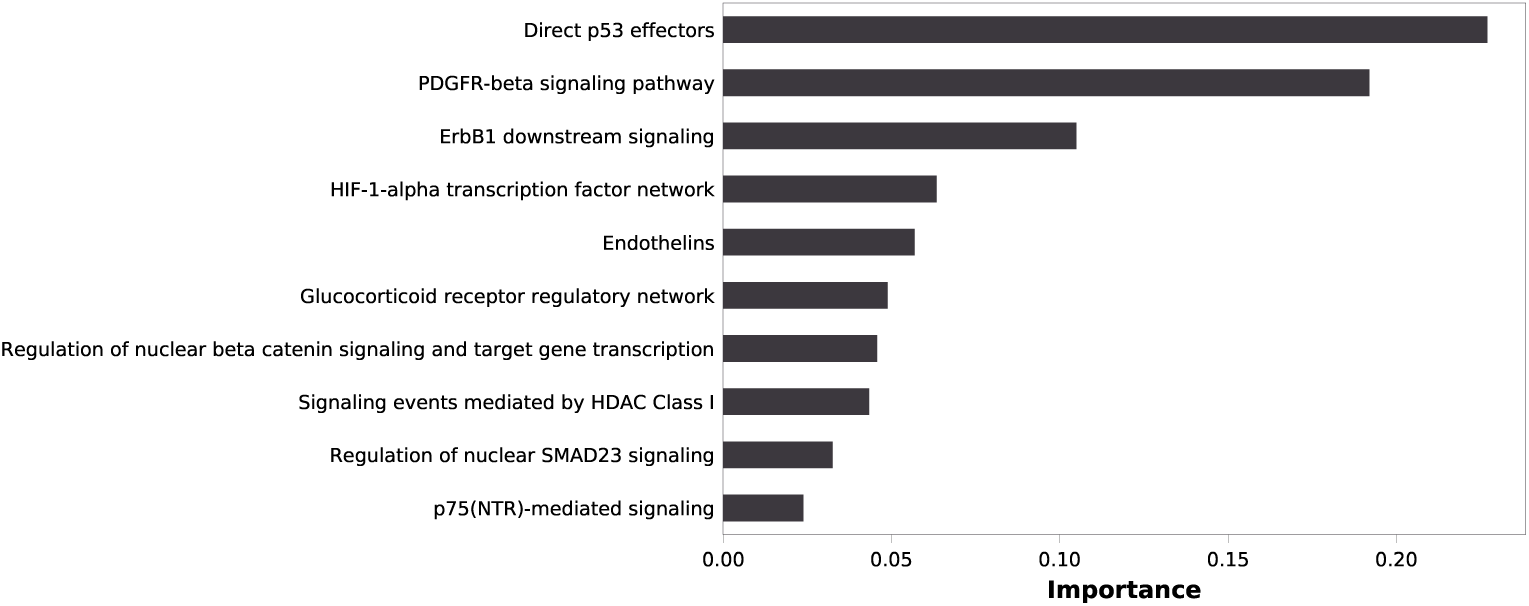
The top 10 pathways, which have the highest relative importance in clustering.

The top relevant pathway in clustering the patients into subgroups emerges as the *the direct p53 effectors pathway* (Figure 5). p53 is a tumor suppressor transcription factor that regulates the cell division to prevent uncontrolled growth of cells [53]. Similarly, the second and the third pathways, *PDGFR-beta signaling* and *ErbB1 downstream signaling pathway* (better known as EGFR) are critical signaling pathways [44]. Additionally, Hypoxia-inducible factors (HIFs) regulates the expressions of many genes that are related to tumorigenesis [1] and [40] shows that HIF1*α* is a target of 14q loss, which is commonly associated with poor prognosis in kidney cancer. The firth pathway is Endothelin pathway and earlier results report that Endotelin-1 promotes cell survival in renal cell carcinoma [32].

## 4 Conclusion and Future Work

We present PAMOGK for discovering subgroups of patients, which not only operates by integrating different omics data sets derived from patients but also incorporates existing knowledge on biological pathways. To corroborate these data sources, we develop a novel graph kernel that evaluates patient similarities based on their molecular alterations in the context of known pathways. We employ a multi-view kernel clustering technique to leverage views constructed by different molecular alteration types and pathways. Our results indicate that the suggested methodology, when applied to KIRC, results in patient clusters that differ significantly in their survival distributions and other clinical parameters. The proposed methodology also provides quantitative evidence for the decisive role of known driver pathways on the clustering process. We further show that PAMOGK performs better compared to the state-of-the-art multi-omics approaches.

One limitation of the current work is that we used the bulk expression results provided by the TCGA project. However, it is known that there could be a high level of intra-tumor heterogeneity [7], and the bulk tumor might include a diverse collection of cells harbouring distinct molecular signatures. Future work would be to adapt PAMOGK framework to single-cell measurements as they become available for large cohorts of patients.

The work can be extended in several directions. In this current work, we use omic datasets containing somatic mutations, gene and protein expression as they give more direct information on the alterations in the pathways. In the future work, one can map the copy number variations and methylation levels as well to the genes and also use them as additional views. Also, we use binary node labels to indicate if the gene for that patient is perturbed or not. PAMOGK can be easily extended to accept continuous node labels to incorporate the extent of the molecular alterations. Furthermore, in the present study, we ignore the direction information of the edges in the pathways. A kernel that explicitly accounts for edge directions can be more devised. In place or addition to the pathways, protein-protein interaction networks can be used to generate views, which will be explored in future work.

## 5 Acknowledgements

This work was supported by the Scientific and Technological Research Council of Turkey (TÜBİTAK) under Grant #117E140. Oznur Tastan thanks to the Science Academy of Turkey under The Young Scientist Award Program (BAGEP). Yasin Tepeli and Ali Burak Unal acknowledge TUBITAK-BIDEB for the 2210-A scholarship program.

## Footnotes

4 The early results of this work are presented as an extended abstract in [49].

5 https://ndexbio.org/#/networkset/8a2d7ee9-1513-11e9-bb6a-0ac135e8bacf

6 https://www.synapse.org/#!Synapse:syn300013

## Notes

https://www.github.com/tastanlab/pamogk

### Competing Interest Statement

The authors have declared no competing interest.

