## Supplementary File for "PAMOGK: A Pathway Graph Kernel based Multi-Omics Clustering Approach for Discovering Cancer Patient Subgroups"

### Supplementary Material for PAMOGK

#### 1 Supplementary Figures

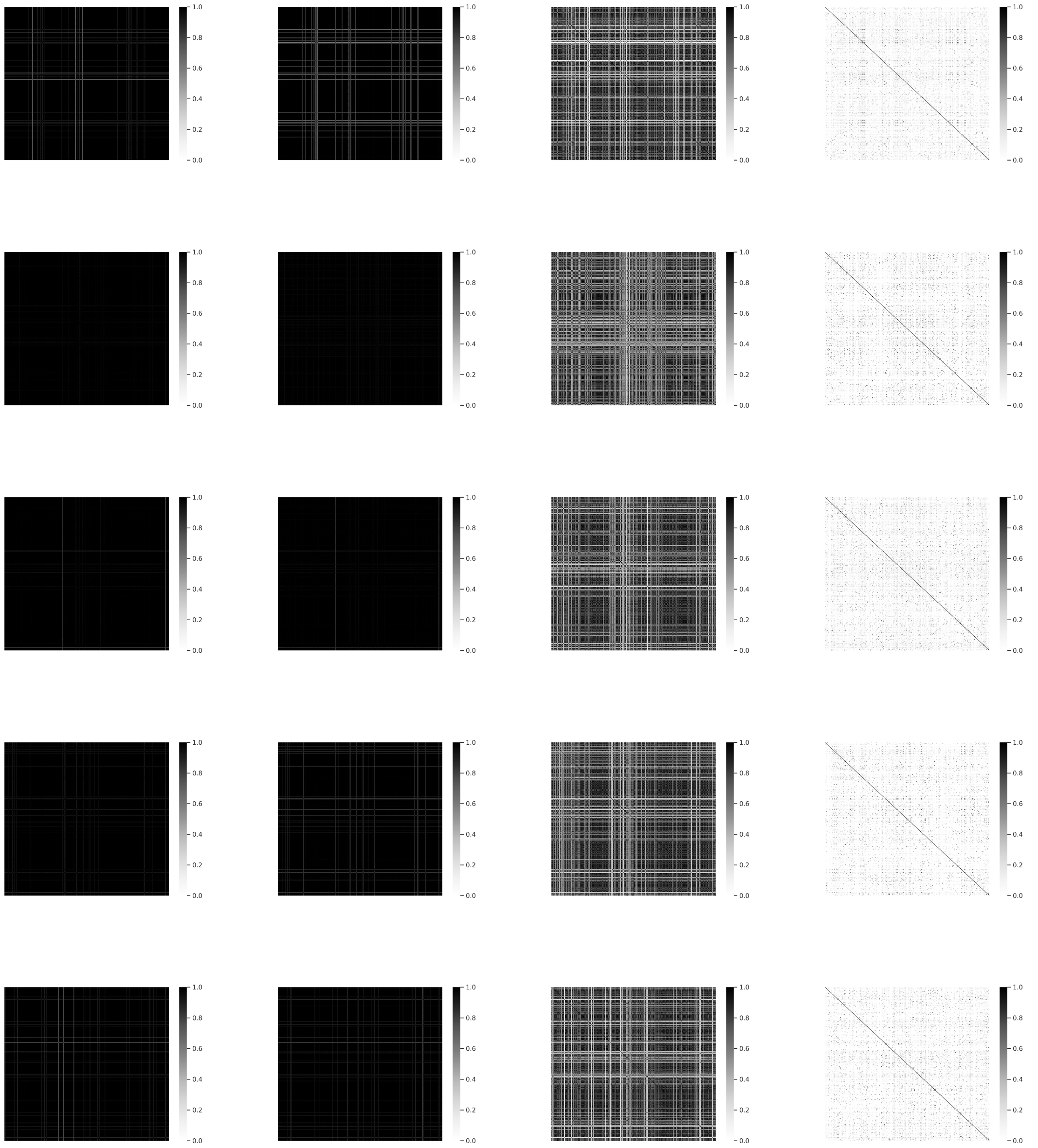

(a) Propagation Kernel (b) Graph Hopper Kernel (c) Wasserstein Weisfeiler Lehman Kernel (d) Smoothed Shortest Path Kernel

Figure 1: Patient-by-patient kernel matrices calculated by different kernel choices. The kernel functions include the propagation kernel, graph hopper kernel, wasserstein weisfeiler lehman, and SmSPK graph kernel methods. Each row corresponds to a randomly chosen pathway and molecular interaction data type. A color black indicates that the two patient similarity is evaluated as 1.

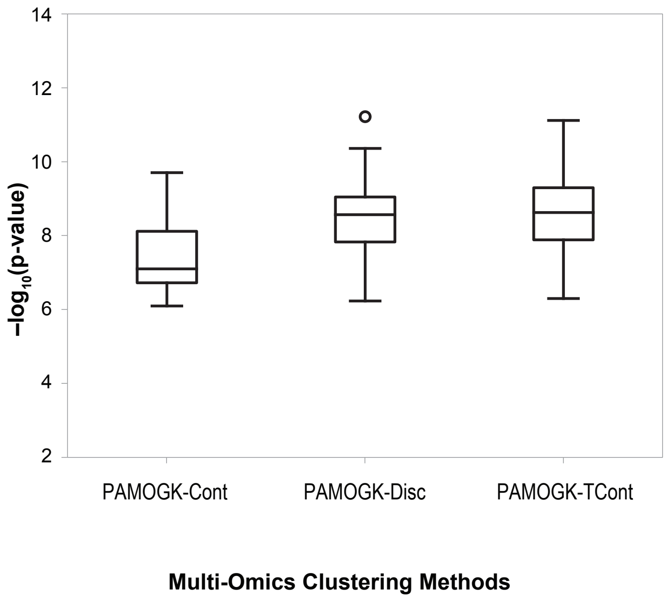


Figure 2: Comparison of different labeling techniques for expression data over 10 different trials. Each trial contains a random subsample of KIRC patients. The boxplot shows the -log ($p$-values) of the log-rank tests conducted on the survival distributions of these clusters. The higher the values, the better the clusters are separated in terms of survival distributions. For the underexpressed and the overexpressed values, we use the z-value as the node attribute. If the z-values are between $-1.96 < z < 1.96$, the node label is assigned to zero(PAMOGK-TCont). In assigning the expression values, we considered different alternative methods, where the node attributes are the z-values (PAMOGK-Cont) or assigned as a binary value where it takes one or zero depending on it is overexpressed or not, and similarly underexpressed or not(PAMOGK-Disc).

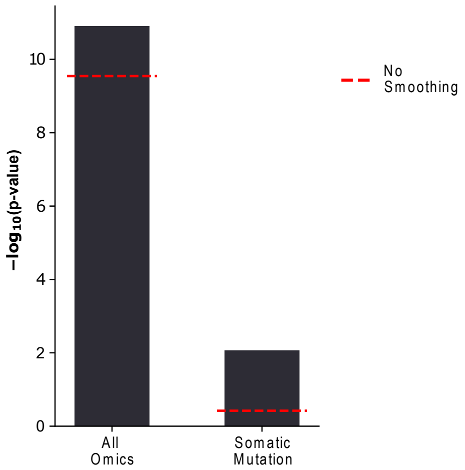


Figure 3: The log-rank test $p$-values obtained for multi-omics data and single-omic data (somatic mutation) with and without smoothing.

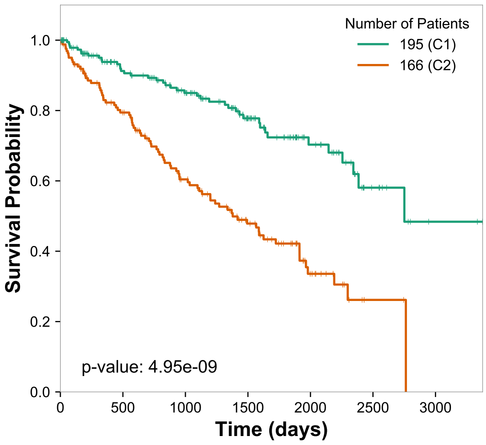


(a) k=2

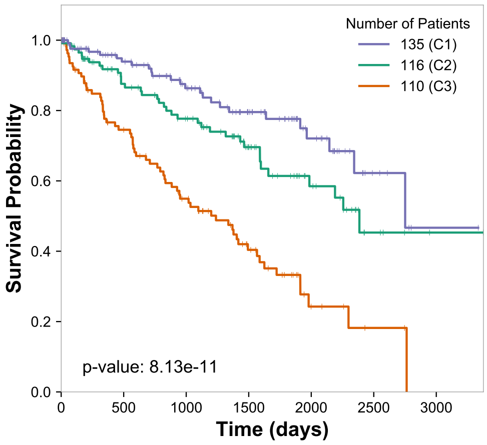


(b) k=3

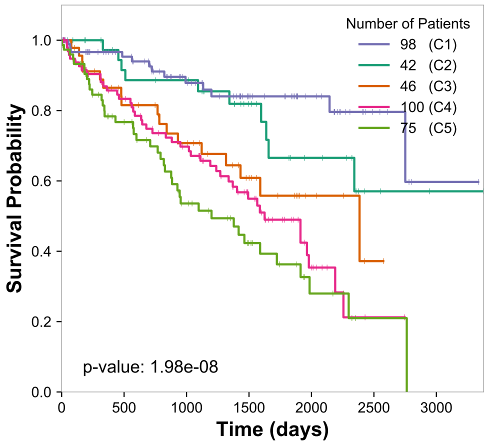


(c) k=5

Figure 4: Kaplan-Meier survival curves of the best clustering solutions for KIRC for different number of clusters k = {2,3,5}. Results obtained with smoothing parameter $\alpha = 0.2$, $\alpha = 0.3$, $\alpha = 0.3$ for k=2(a), k=3(b), k=5(c), respectively. The p-value was obtained from a log-rank test between the groups.

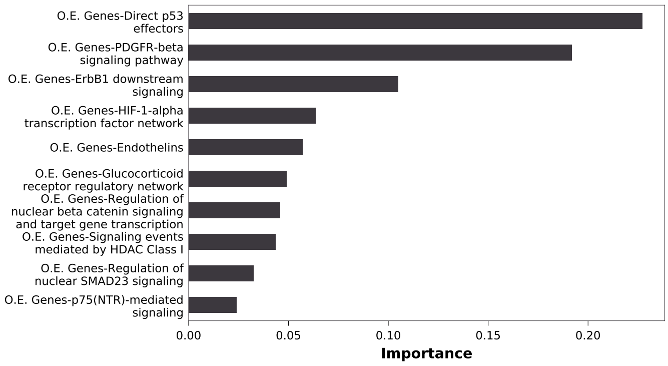


Figure 5: Top 10 most influential pathway-alteration type pairs. O.E. stands for overexpressed and U.E. stands for under-expression. The relative importance is calculated based on the weights assigned to each kernel matrix of the associated pair by the MKKM-MR algorithm. The results are obtained for the best clustering solution, where the number of cluster is 4, kernel matrices are calculated using SmSPK with smoothing parameter $\alpha = 0.3$.

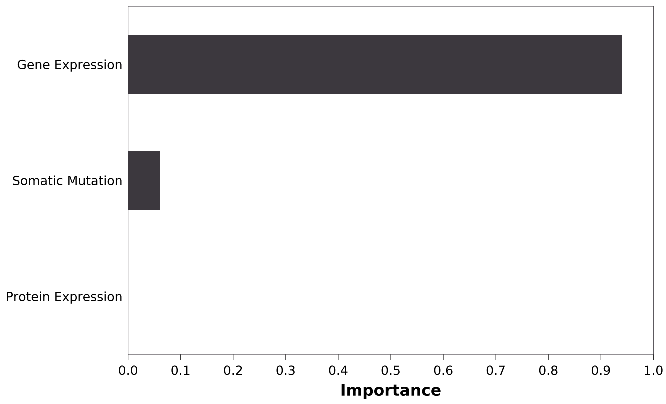


Figure 6: Relative importance of the three omic data types. One data type weight was calculated by summing up the kernel weights that is available for molecular alteration type and pathway pair. The results are obtained for the best clustering solution, where the number of clusters is 4, and the kernel matrices are calculated by SmSPK with smoothing parameter $\alpha = 0.3$.

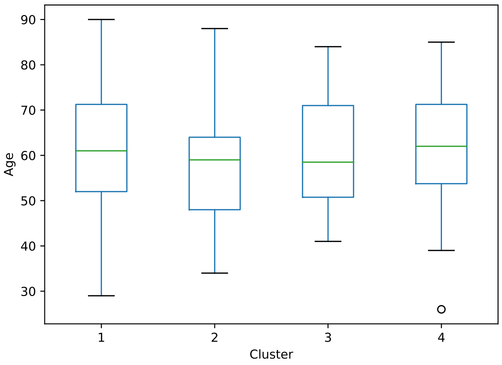


Figure 7: Age distribution of patients in each identified RCC cluster. No statistical significance across groups is detected via one-Way ANOVA test ( p-value = 0.143)

### 2 Supplementary Tables

Table 1: Data sources and their download dates of datasets used in PAMOGK experiments.

| Data | Source | Download Date |
|---|---|---|
| **Somatic Mutations** | https://www.synapse.org/#!Synapse:syn1701259 | March 24, 2019 |
| **Gene Expression** | https://www.synapse.org/#!Synapse:syn417925.5 | April 24, 2019 |
| **Protein Expression** | https://www.synapse.org/#!Synapse:syn416783.3 | April 24, 2019 |
| **Clinical Data** | https://www.synapse.org/#!Synapse:syn417024.7 | April 24, 2019 |
| **Pathway Data** | https://ndexbio.org/#/networkset/8a2d7ee9-1513-11e9-bb6a-0ac135e8bacf | April 24, 2019 |

Table 2: Statistics of 165 pathways

| | Average | Median | Std. Dev. | Max. number of | Min. number of |
|---|---|---|---|---|---|
| **Nodes** | 44.648 | 42 | 23.952 | 142 | 2 |
| **Edges** | 231.994 | 181 | 198.078 | 1277 | 1 |

Table 3: Hyperparameters used in different algorithms. RBF values are selected using the median heuristic.

| Parameter | Symbol | Used in | Possible value(s) |
|---|---|---|---|
| **Number of clusters** | $\boldsymbol{k}$ | All clustering methods | $\{2, 3, 4, 5\}$ |
| **Smoothing** | $\boldsymbol{\alpha}$ | Kernel construction of SmSPK | $\{0, 0.01, 0.05, 0.1, 0.2, 0.3, 0.4, 0.5, 0.6, 0.7, 0.8, 0.9\}$ |
| **Trade-off** | $\boldsymbol{\lambda}$ | MKKM-MR | $2^{\{-15,-12,-9,\cdots,9,12,15\}}$ |
| | | Somatic Mutation | 6.41e-03 |
| **RBF** | $\gamma$ | Gene Expression | 8.01e-04 |
| | | Protein Expression | 1.11e-01 |
| **Number of neighbors** | **Ks** | SNF | 20 |
| **Number of iterations** | **Ts** | SNF | 20 |
| **Max. number of iterations** | **Tl** | LMKKM | 50 |

### 3 Statistical Association of the KIRC Clusters with the Clinical Parameters

Table 4: Summary of TNM staging according to AJCC [1].

| Tumor | Definition |
|---|---|
| **T1** | Tumor $\leq 7$ cm in greatest dimension, limited to the kidney |
| **T2** | Tumor $\geq 7$ cm in greatest dimension, limited to the kidney |
| **T3** | Tumor extends into major veins or perinephric tissues but not into the ipsilateral adrenal gland and not beyond Gerota's fascia |
| **T4** | Tumor invades beyond Gerota's fascia |
| **M0** | No distant metastasis |
| **M1** | Distant metastasis |
| **Stage I** | T1 - M0 |
| **Stage II** | T2 - M0 |
| **Stage III** | T1 or T2 - M0(Additionally metastasis in lymph node)<br>T3 - M0 |
| **Stage IV** | T4 - M1<br>Any T - M0 |
| **GX** | Grade cannot be assessed - Tumor cell and tissue is close to normal |
| **G1** | Well differentiated - Tends to grow slowly |
| **G2** | Moderately differentiated - Tends to grow rapidly and faster |
| **G3** | Poorly differentiated - Tends to grow rapidly and faster |
| **G4** | Undifferentiated |

Note that the Cluster 1 is the patient subgroup with the best prognosis and the Cluster 4 is the worst prognosis. Refer to Figure 3a in main text for the cluster ids and their survival distributions and please refer to Table 4 for definition of clinical terms.

Table 5: Contingency table for gender vs clusters. The chi-squared test results in $\chi^2 = 2.893$, $p = 0.408$, df= 3

| **Gender** | **Female** | **Male** | **All** |
|---|---|---|---|
| **Cluster No** | | | |
| **1** | 39 | 63 | 102 |
| **2** | 22 | 40 | 62 |
| **3** | 26 | 56 | 82 |
| **4** | 32 | 83 | 115 |
| **ALL** | 119 | 242 | 361 |

Table 6: Contingency table for tumor stage vs clusters. The chi-squared test results in $\chi^2 = 52.603$, $p = 3.476e - 08$, df $= 9$

| **Stage** | **I** | **II** | **III** | **IV** | **All** |
|---|---|---|---|---|---|
| **Cluster No** | | | | | |
| **1** | 55 | 12 | 25 | 10 | 102 |
| **2** | 39 | 5 | 10 | 8 | 62 |
| **3** | 50 | 3 | 18 | 11 | 82 |
| **4** | 24 | 13 | 44 | 34 | 115 |
| **ALL** | 168 | 33 | 97 | 63 | 361 |

Table 7: Contingency table for primary tumor pathological spread vs cluster. Chi-squared test results in $\chi^2 = 49.479$, $p = 1.349e - 07$, df $= 9$

| **Pathologic Spread** | **T1** | **T2** | **T3** | **T4** | **All** |
|---|---|---|---|---|---|
| **Cluster No** | | | | | |
| **1** | 57 | 14 | 30 | 1 | 102 |
| **2** | 40 | 6 | 16 | 0 | 62 |
| **3** | 50 | 5 | 26 | 1 | 82 |
| **4** | 26 | 16 | 69 | 4 | 115 |
| **ALL** | 173 | 41 | 141 | 6 | 361 |

Table 8: Contingency table of distant metastasis pathological spread vs cluster. The chi-squared test results in $\chi^2 = 18.327$, $p = 3.766e - 04$, df = 3

| **Pathologic Spread** | **M0** | **M1** | **All** |
|---|---|---|---|
| **Cluster No** | | | |
| **1** | 93 | 9 | 102 |
| **2** | 54 | 8 | 62 |
| **3** | 70 | 12 | 82 |
| **4** | 81 | 34 | 115 |
| **ALL** | 298 | 63 | 361 |

Table 9: Contingency table for neoplasm histological grade vs clusters. The chi-squared test results in $\chi^2 = 65.608$, $p = 2.104e - 09$, df = 12

| **Histologic grade** | **G1** | **G2** | **G3** | **G4** | **GX** | **All** |
|---|---|---|---|---|---|---|
| **Cluster No** | | | | | | |
| **1** | 2 | 61 | 33 | 6 | 0 | 102 |
| **2** | 2 | 26 | 26 | 8 | 0 | 62 |
| **3** | 1 | 35 | 38 | 8 | 0 | 82 |
| **4** | 0 | 21 | 52 | 41 | 1 | 115 |
| **ALL** | 5 | 143 | 149 | 63 | 1 | 361 |
